# A searchable metadata network graph for microbiome metabolomics

**DOI:** 10.64898/2026.02.04.703849

**Authors:** Vincent Charron-Lamoureux, Shipei Xing, Abubaker Patan, Corinn Walker, Ricardo-Almada Monter, Yasin El Abiead, Haoqi Nina Zhao, Lucas Patel, Yuhan Weng, Antonio Gonzalez, Gail Ackermann, Victoria Deleray, Vishant Gandhi, Ipsita Mohanty, Andres Mauricio Caraballo-Rodriguez, Kine Eide Kvitne, Simone Zuffa, Annabelle Norman, Anthony Martin, Loryn Chin, Rocio Paz-Gonzalez, Marta Sala-Climent, Nadia Suryawinata, Jasmine Zemlin, Harsha Gouda, Zhewen Hu, Grant Norton, Prajit Rajkumar, Anthony JA Molina, Jaclyn Bergstrom, Monae Pinner, Sadie Giddings, Allegra T. Aron, Liang Liang, Samira Dahesh, Santosh Lamichhane, Erin R. Reilly, Victor Nizet, Anna Skrip, April L. Lukowski, Selene F.H. Shore, Subhomitra Ghoshal, Melinda A. Engevik, Thomas D. Horvath, Simone Renwick, Julius Agongo, Maria L. Marco, Sarkis K. Mazmanian, Mingxun Wang, Heejung Yang, Daniel McDonald, Monica Guma, Evi Stegmann, Naybel Hernandez Perez, Paolo Stincone, Eric Kemen, Abzer Kelminal Pakkir Shah, Lars Bode, Daniel Petras, Dionicio Siegel, Manuela Raffatellu, Andrew D. Patterson, Suzanne Devkota, Adrian Jinich, Rob Knight, Karsten Zengler, Pieter C. Dorrestein

## Abstract

Establishing the biological context of microbial metabolites remains a major challenge. We present microbiomeMASST, a metadata-driven network graph that maps metabolites across 467 available datasets with 144,424 mass spectrometry files from humans, animals, and microbial culture systems. MicrobiomeMASST integrates monocultures, synthetic communities, and host-associated samples across multiple body sites and plants. MS/MS spectra can be queried to trace occurrence across hosts, experimental conditions, and interventions, enabling cross-study integration. We demonstrate this framework by contextualizing microbial-conjugated bile acids and interrogating microbiome-mediated drug metabolism. Screening gut bacteria revealed deprolylation of the angiotensin-converting enzyme (ACE) inhibitor prodrug enalapril. Using microbiomeMASST, we traced this metabolite across human cohorts, microbial isolates, environmental samples, and in *Gorilla gorilla*. Structural modeling and enzymatic assays showed that microbial deprolylation abolishes ACE inhibition, thereby inactivating its therapeutic effect. Together, microbiomeMASST links MS/MS spectra to biological context, converting isolated observations into an interpretable microbiome map for cross-study analysis.

## Main

Microbiomes shape the host and its surrounding ecosystem, mediated through metabolites that can be derived from the host, medications^1–3^, diets^4^, pesticides^5^, or other environmental exposures^6^. Microbiota can influence disease risk^7,8^, shaping health trajectories across life stages^9,10^ and be influenced by lifestyle^11^, often with effects extending far beyond their immediate microbial environment^12^. Although crucial across biological systems, microbial metabolites are frequently reported in individual studies, and often studied in individual biological matrices such as feces, urine, serum, saliva, or plasma^13,14^. Once a newly discovered microbial metabolite is described, it is often published solely within the context of its initial observation, without broader insight into its larger biological and ecological context^15,16^. This limited perspective leaves many questions unanswered: Is the metabolite relevant across different health conditions? How do medications or diets modulate its levels? Can it reach distant organs such as the brain or eyes? How do factors such as diet, gut pH, or anatomical location along the digestive tract shape its abundance? Are individual microbes sufficient for its production, or is a community required? Is it produced under monocolonization conditions in animal models? Does its presence vary between healthy and diseased states? Does it accumulate systemically, and if so, in which organs? If first discovered in animals, does it translate to humans, and at what stage of life: early after birth, during development, old age, or even after death? How do diurnal or circadian rhythms affect its levels? And how do the abundances of producing microbes shift across microbiome-relevant conditions? While many of these questions are most immediately framed in human health contexts, similar challenges apply across animals, plants, and environment, as well as bacterial monocultures and synthetic communities. Answering these questions requires dedicated human or animal cohorts, substantial funding, years of systematic study, and resources beyond the reach of most individual laboratories.

Currently a growing resource of publicly accessible untargeted metabolomics datasets exists that could, in principle, help answer many of these questions^17,18^. However, practical barriers limit its broader use: cross-study analysis requires downloading, processing of raw data, harmonizing metadata, and performing study-by-study interpretation, all of which require substantial time and expertise to navigate. Some sample metadata in public projects may be defined using structured ontologies, such as taxonomy or tissue type, enabling data-driven analyses, as demonstrated with existing tools like microbeMASST^19^, tissueMASST^20^, or via PanReDU metadata harmonization using controlled vocabularies (e.g., age or sex)^21^. Yet most experimental details such as dietary interventions, oxygen or carbon dioxide levels, drug treatments, microbial colonization experiments, surgical procedures, sample collection times, and provision of individual compounds are not fully captured by existing ontologies and become effectively lost in ontology- or controlled-vocabulary-restricted repository scale searches.

Network graphs offer a complementary framework to ontology-restricted metadata by representing relationships among metabolites, sample types, experimental conditions, and studies, thereby enabling cross-study contextualization and hypothesis generation. In data science, a network graph represents entities (nodes) and their relationships (edges), allowing connections to be drawn across otherwise disparate datasets. Unlike established ontology-only systems, network graphs can also integrate heterogeneous data types - including non-standardized sample information such as experimental conditions - enabling discovery of higher-order patterns, identification of clusters, and mapping of relationships across studies that would otherwise remain siloed^22–24^. When no standardized ontologies exist, as is the case for many experimental conditions, metadata connections must be curated in a computer-readable format one by one and file by file. This manual curation enables downstream computational integration and analysis. In metabolomics, such graphs can reveal how microbial metabolites and their producers are connected across health states, diets, model systems, and interventions.

To demonstrate this principle for microbiome applications, we built a network graph linking study metadata with corresponding liquid-chromatography tandem mass spectrometry (LC-MS/MS) data files, with emphasis on microbiome-relevant human, animal, and microbial culturing conditions. This approach allows questions to be answered across contexts rather than within a single study. In total, our network graph spans 467 studies, resulting in 1,596 different metadata entries that make up the edges of the graph by going through the original papers, reaching out to original depositors of the data to get additional details when needed and then building connections to form the graph.

To further empower the scientific community to contextualize microbial molecules, each node in the network graph is linked back to the raw MS/MS data via a Universal Spectrum Identifier (USI)^25^. The USI provides a unique, reproducible reference for any MS/MS spectrum, and the USI resolver ensures that it can be traced back to the exact raw spectrum in its original repository or data source. This linkage bridges into the GNPS2 fast MASST (FASST) ecosystem^26,27^, allowing researchers to input the MS/MS spectrum of a microbiome-derived molecule and immediately query it across the studies that are part of this network graph. Instead of a list of matches, results are embedded in an interpretable, network graph that highlights the distribution of the molecule across studies, conditions, hosts, and microbial systems.

This framework transforms isolated discoveries of microbial metabolites into a broader, contextual microbiome map, allowing users to see not only where a molecule occurs in a single study, but also where it appears and under what experimental conditions across diets, medications, microbiome manipulations, or host species. This contextualization is even feasible when the molecule itself was not annotated when the data was originally deposited. By integrating metadata and experimental context into the graph structure, researchers can leverage single-study observations and contextualize how a metabolite fits into the broader ecology of the microbiome.

## Results

Establishing microbiome-relevant context for MS/MS spectral searches required a systematic evaluation of both data availability and associated metadata (sample information) across public repositories (**Fig. 1a**). We included datasets from the metabolomics repositories GNPS/MassIVE^24^, MetaboLights^28^, Metabolomics Workbench^29^ (**Supplementary Table S1**). In cases where metadata were insufficient or ambiguous but the underlying data appeared relevant to microbiome science, we enhanced the metadata through direct communication with the depositing authors or authors involved in the original studies.

**Fig. 1.**
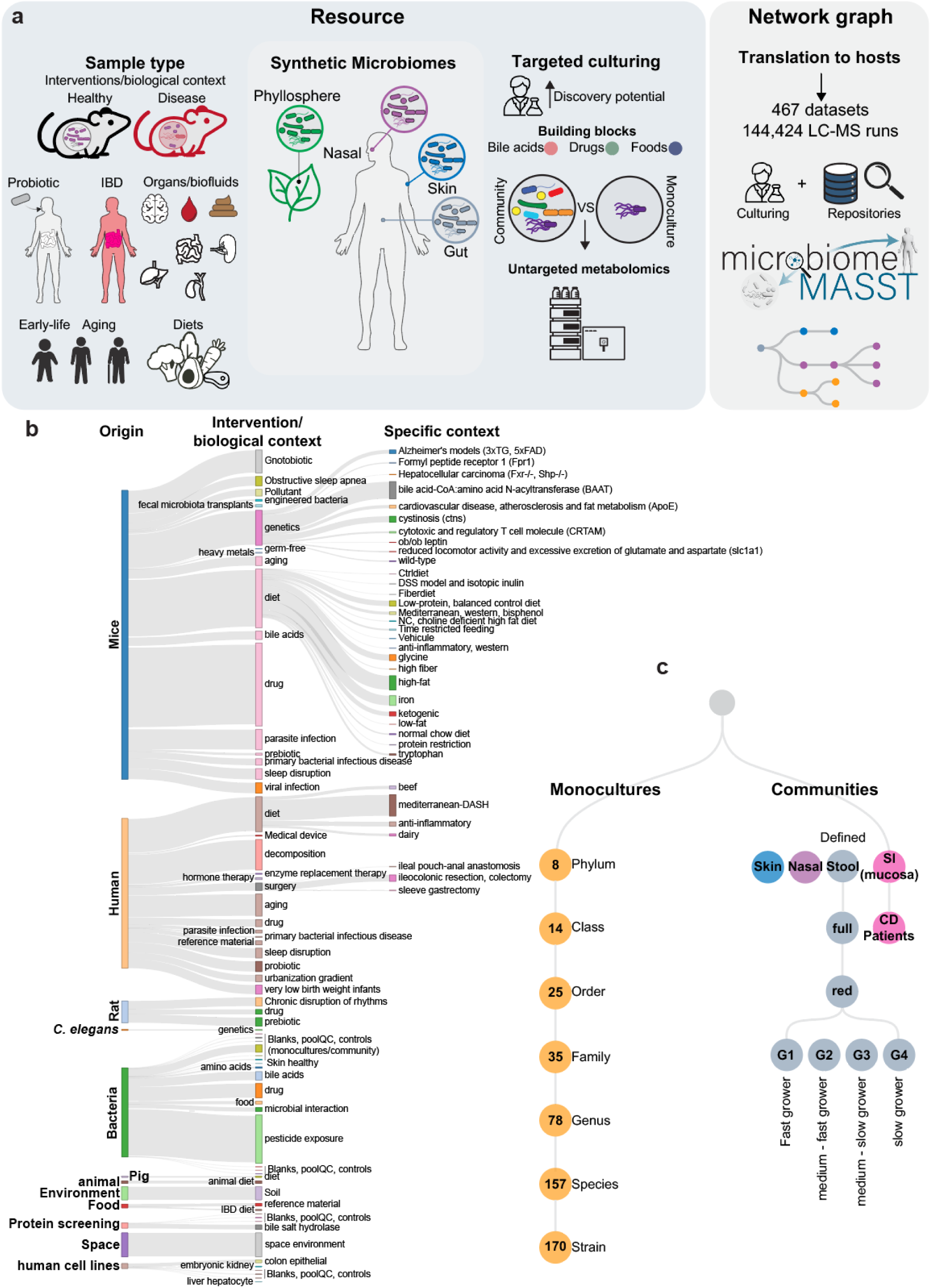
Creation of a microbiome-focused network graph to translate microbial metabolites into host biology. **a)** The network graph is based on biological interventions and contexts mined from metabolomics repositories (GNPS/MassIVE; 274 studies), (Metabolights; 86 studies), (Metabolomics Workbench; 107 studies). Monocultures and communities were profiled using targeted culturing approaches to increase the discovery potential of microbial metabolites by supplementing to the culture medium with building blocks relevant for human health (e.g., bile acids, drugs, and foods) prior to untargeted metabolomics analysis. This network graph is embedded into the search engine microbiomeMASST. **b)** Overview of a portion and one layer of the network graph-based microbiomeMASST and its organization, showcasing genetic mouse models, human, mice diets, and surgical procedures. The width of the bars is proportional to the number of files available for each category; for example, mouse drug interventions comprise 3,822 files, mouse diets 3,563 files, human diets 1812 files, and human aging data 1201 files. **c)** MicrobiomeMASST provides strain-level resolution, encompassing 170 strains spanning 157 species, 78 genera, 35 families, 25 orders, 14 classes, and 8 phyla. The resource includes data from monocultures, defined and sub-communities, and dysbiotic communities derived from the small intestinal mucosa of Crohn’s disease patients. Sample types represented include nasal, stool, skin, and plant-associated (phyllosphere) data. IBD, inflammatory bowel disease; SI, small intestine; CD, Crohn’s disease.

Metadata attributes span both interventional and observational study designs and include colonization status (e.g., germ-free, monocolonized mice, specific-pathogen free), dietary interventions (e.g., high-/low-fat, high-/low-protein, time-restricted feeding), molecule exposures (e.g., medications, cholesterol, bile acids^30^), life stage (newborns, aging^9,10^, postmortem decomposition^31^), diurnal/circadian patterns, sex, genetic knockouts^32^, sampling strategies (including ingestible gastrointestinal tract pill sampling devices^33^), organs and biofluids, microbiome-relevant health states (e.g., inflammatory bowel disease^34,35^, diabetes, Alzheimer’s disease) and surgical procedures^35^.

To build a cross-species graph, we integrated curated datasets across humans (32 body parts, biofluids, and cell types), rodents (65 biospecimen types including stool, blood, and urine), and pigs (27 sites), enabling assessment of whether microbial metabolites appear systemically across different mammals. Beyond host biology, we also curated microbial culture experiments spanning multiple levels of complexity: monocultures at the strain level (170 strains belonging to 157 species, 78 genera, 35 families, 25 orders, 14 classes, 8 phyla), defined synthetic communities derived from different sample types (nasal, skin, stool) and health states (Crohn’s disease, Alzheimer’s disease, diabetes), and artificial gut systems with dynamic pH conditions^36^. The phyllosphere was included as a well-characterized plant-associated microbiome, represented by a synthetic microbial community derived from *Arabidopsis thaliana* phyllosphere, for which an untargeted LC-MS/MS dataset with metadata was available. The network graph in microbiomeMASST captures detections of microbial cultures in community contexts, which is important because some metabolites are produced through microbial interactions rather than in axenic culture. Experimental contexts within microbiomeMASST include microbial incubations with hundreds of health-relevant molecules (e.g., bile acids, fatty acids^12,37^, polyamines), 164 commonly prescribed drugs^3,38^ (e.g., metoprolol, escitalopram), 268 food molecules and representative foods (e.g., apple, cow milk)^39^, and enzyme assays (currently 126 bile salt hydrolases/transferases primarily from gut microbes, with ongoing expansion planned; **Figs. 1b–c, Supplementary Tables S2–S6**).

To aid in the data science workflow and visualization, metadata fields were harmonized through manual curation using scripted transformation in RStudio^40^, in which synonymous terms (e.g., “stool,” “fecal,” “feces”) were collapsed into a single controlled term and aligned with the USI and dataset identifier framework so that all datasets could be accessed via the USI resolver. Using these harmonized datasets, we constructed a microbiome-centric network graph in which each metadata attribute is represented as a node linked by edges according to biological or experimental context. This graph was iteratively refined through visualization and expert interpretation, with the initial layers capturing sample origin, intervention or biological context, and specific experimental attributes. Sankey diagrams illustrate representative flows of sample information (**Figs. 1b-c**). Importantly, individual datasets could map to multiple interpretable units - for example, a mouse study involving both diet manipulation (high-fat vs. normal chow) and antibiotic treatment was connected to both the diet and drug intervention categories.

Altogether, version 1 of the microbiome network graph integrates 467 studies that are in public metabolomics data repositories encompassing 144,424 LC-MS/MS runs and more than 278 million MS/MS spectra (most collected in positive ionization mode), spanning human, animal, microbial, dietary, drugs, and built-environment microbiomes, from rural households to the International Space Station. The graph comprises 1,596 nodes (median 6 LC-MS/MS files per intervention type, interquartile range, 3-99; range 1-6078 files) and 1,595 edges, and is now deployed in the GNPS2/fastMASST (FASST) ecosystem. A dedicated web interface allows users to input MS/MS spectra from a USI or manual upload, returning contextualized matches within user-defined scoring thresholds visualized as a network graph https://masst.gnps2.org/microbiomemasst/ (**Supplementary Figure 1**). This framework enables hypothesis generation by connecting metabolite features to their microbial producers, host distribution, health associations, or environmental origins, as illustrated in the following bile acid examples.

### Microbially-conjugated bile acids: contextualized insights

MicrobiomeMASST enables contextualization of microbial metabolites across studies and to illustrate its utility, we highlighted representative short vignettes, each accompanied by videos that demonstrate how to use the network graph and the depth of contextualization it can provide. For illustration, we focus on diverse and complex microbially conjugated bile acids, some of which are described here for the first time. We then show how microbiomeMASST provides biological context to these molecular discoveries.

Microbiome-mediated bile acid transformations generate a diverse array of microbially conjugated bile acids increasingly recognized for their biological significance^41^. Polyamine conjugates such as N-acetyl spermidine and cadaverine with bile acids have been detected in human and animal samples by our group, including in lion feces and human stools, but their microbial producers had not been established because no matches were obtained from the 60,781 taxonomically curated LC-MS/MS files of monocultures^42^. It was not clear that microbes could synthesize these compounds. Using microbiomeMASST, we confirmed and contextualized two reported polyamine conjugates, trihydroxylated bile acid–N-acetyl spermidine and trihydroxylated bile acid–cadaverine, produced specifically by *Thomasclavelia spiroformis* in monoculture and in pooled microbial communities (**Figs. 2a-b; <u>Video 1</u>**). This identification was possible because the microbiome network graph contains datasets in which polyamines were added to the growth media, providing the first evidence to support the hypothesis that these polyamine–bile acid conjugates can be synthesized by bacteria. Both conjugates were observed in digestive tract-associated samples and in breastmilk, and they were detected across multiple biological contexts, including human and mouse datasets involving dietary interventions, prebiotic treatments, and pathological states such as liver cancer models. Typically, this level of contextualization would require time-consuming examination of one study at a time. Similarly, we observed that deoxycholyl–2,3-diaminopropionic acid, another previously unreported diamine conjugate, was produced by *Coprococcus eutactus* in culture and independently detected in a purified bile salt hydrolase assay spiked with substrates that included 2,3-diaminopropionic acid, supporting a microbial and microbial enzyme-mediated origin for this bile acid (**Supplementary Figure 2; <u>Video 2</u>**). Analysis of *Coprococcus eutactus* cultures using LC-ion-mobility MS, with retention and drift time matching the synthetic standard, confirmed microbial production and reinforced the power of microbiomeMASST to link chemical structure, microbial producers, and experimental context (**Supplementary Figure 2e)**.

**Fig. 2.**
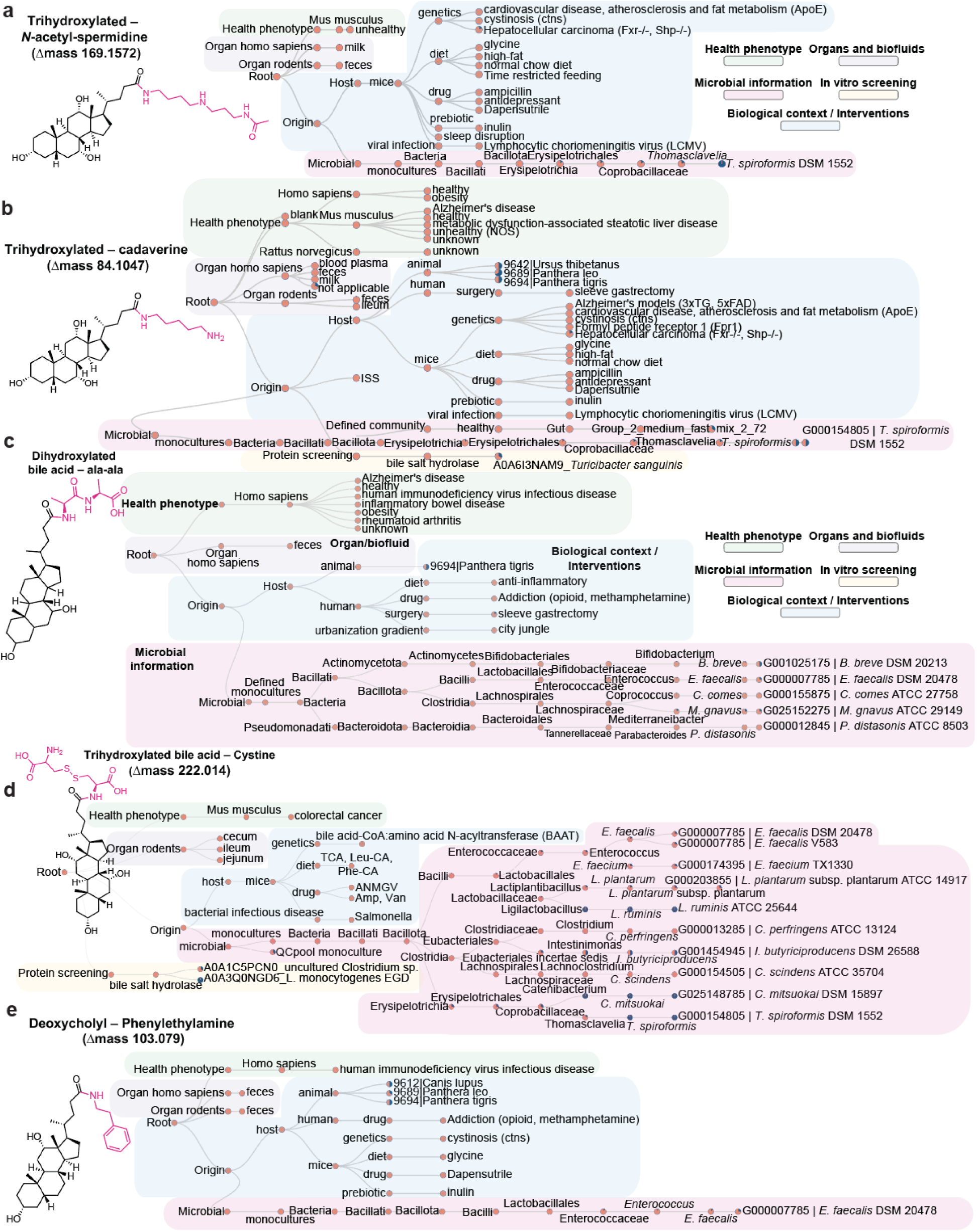
Contextualization of known and unknown bile acids using the microbiomeMASST network graph. **a)** MicrobiomeMASST results for trihydroxylated bile acid–N-acetyl-spermidine (library ID CCMSLIB00010221653). **b)** Trihydroxylated bile acid–cadaverine (library ID CCMSLIB00010224778). **c)** Dihydroxylated bile acid–alanine-alanine (library ID CCMSLIB00015103143). **d)** Cholyl–cystine (library ID CCMSLIB00010220367). **e)** Deoxycholyl–phenylethylamine (library ID CCMSLIB00015466534). Each panel shows detections across bacterial monocultures (pink), synthetic communities (pink), diverse biological contexts and interventions (blue), organs and biofluids (purple), and health phenotype (green). The MS/MS spectral similarities and retention time matching of the structures presented in d and e can be found in **Supplementary Figure 3c,d**.

Another set of conjugated bile acids involves dipeptide amidates. Lithocholyl–ala-ala and deoxycholyl–ala-ala, alanine–alanine conjugates of lithocholic acid and deoxycholic acid, were recently described in humans and dogs through a mass spectrometry-based bile acid identification study, but microbial producers had not been identified (**Fig. 2c; <u>Video 3</u>**)^43^. These dipeptide conjugates were observed with lithocholic acid, deoxycholic acid, and chenodeoxycholic acid, but not with cholic acid, despite cholic acid being more abundant and a precursor to these secondary bile acids. The dipeptide conjugate was synthesized and further validated via targeted synthesis and MS/MS spectral matching, detected in humans and selected animal models, and observed across diverse health contexts including inflammatory bowel disease (IBD), Alzheimer’s disease, and aging (**Supplementary Figure 3a,b**). Microbial cultures from six different strains spanning three taxonomic classes–Actinomycetes, Bacilli, and Clostridia–also produced this conjugate.

Additional conjugates were observed during microbial culture experiments for the first time. This included cholyl–cystine, a trihydroxylated bile acid conjugated to cystine, validated using a synthetic standard in culture of *T. spiroformis* (**Fig. 2d; Supplementary Figure 3c; <u>Video</u> <u>4</u>**). Using microbiomeMASST, this metabolite was also detected in mouse small intestine, cecum, and proximal large intestine, and may form via direct enzymatic conjugation of cystine or through oxidation of cysteine in culture media (**Fig. 2d**). We also detected and validated deoxycholyl–phenylethylamine (**Supplementary Figure 3d**), produced by *E. faecalis*. In this case, phenylethylamine was not added to the culture media but is a known endogenous product of *E. faecalis* metabolism^44^. Using microbiomeMASST, the MS/MS of this molecule could be traced into humans, tigers, and mice across diverse contexts including diets, drug treatments, and mouse studies with genetic modifications (e.g., knockouts) (**Figs. 2e ; <u>Video 5</u>)**. Collectively, these representative vignettes offer broader insight into the microbial production of bile acid conjugates and illustrate how systematic cross-study mapping contextualizes their occurrence across hosts, health states, and experimental interventions.

### Human gut microbes metabolize the angiotensin-converting enzyme inhibitor prodrug enalapril

Next we leveraged our microbiome-centric network graph to interrogate the microbial metabolism of drugs, an area in which gut microbes have been shown to influence treatment outcomes in Parkinson’s disease^45^, IBD^8^, and cancer chemotherapy^46^. *In vitro* screens show that gut bacteria can deplete, biotransform, and accumulate a wide range of medications, potentially altering drug efficacy and toxicity^1,2,47^. Within microbiomeMASST, we included a screen of 164 commonly prescribed drugs incubated with gut-derived synthetic communities. As an illustrative example, we queried enalapril, an antihypertensive prodrug used in our drug screening assay, to illustrate how the graph structure is interpreted **(Supplementary Figure 4a-b)**. In humans, enalapril is converted by hepatic esterases into enalaprilat–its active form^48^.

Despite hundreds of biotransformations reported in high-throughput studies, only a handful of microbial-derived drug metabolites have been validated *in vivo*^1,2^, creating a major translational bottleneck. To address this, we used microbiomeMASST’s cross-study analyses capabilities to trace a drug metabolite first observed when cultured with synthetic microbial communities (SynComs) from the gut *in vitro*. In a large culturing drug screen experiment, we found an unannotated feature (*m/z* 280.154; RT 5.28 min) that showed a significant increase across all four groups of bacteria (fast growers, medium-fast growers, medium-slow growers, and slow growers) compared to the media control after 72 h **(Fig. 3a)**. Feature-based molecular networking^23^ revealed that this feature clustered with enalapril and exhibited a Δ *m/z* −97.05, consistent with loss of proline **(Fig. 3b)**. Therefore, we hypothesized that gut microbes might cleave the proline moiety, converting enalapril to desprolyl-enalapril. MS/MS spectral matching and retention time comparison between the authentic standard and biological sample confirmed the formation of desprolyl-enalapril produced by human gut communities **(Fig. 3c)**.

**Fig. 3.**
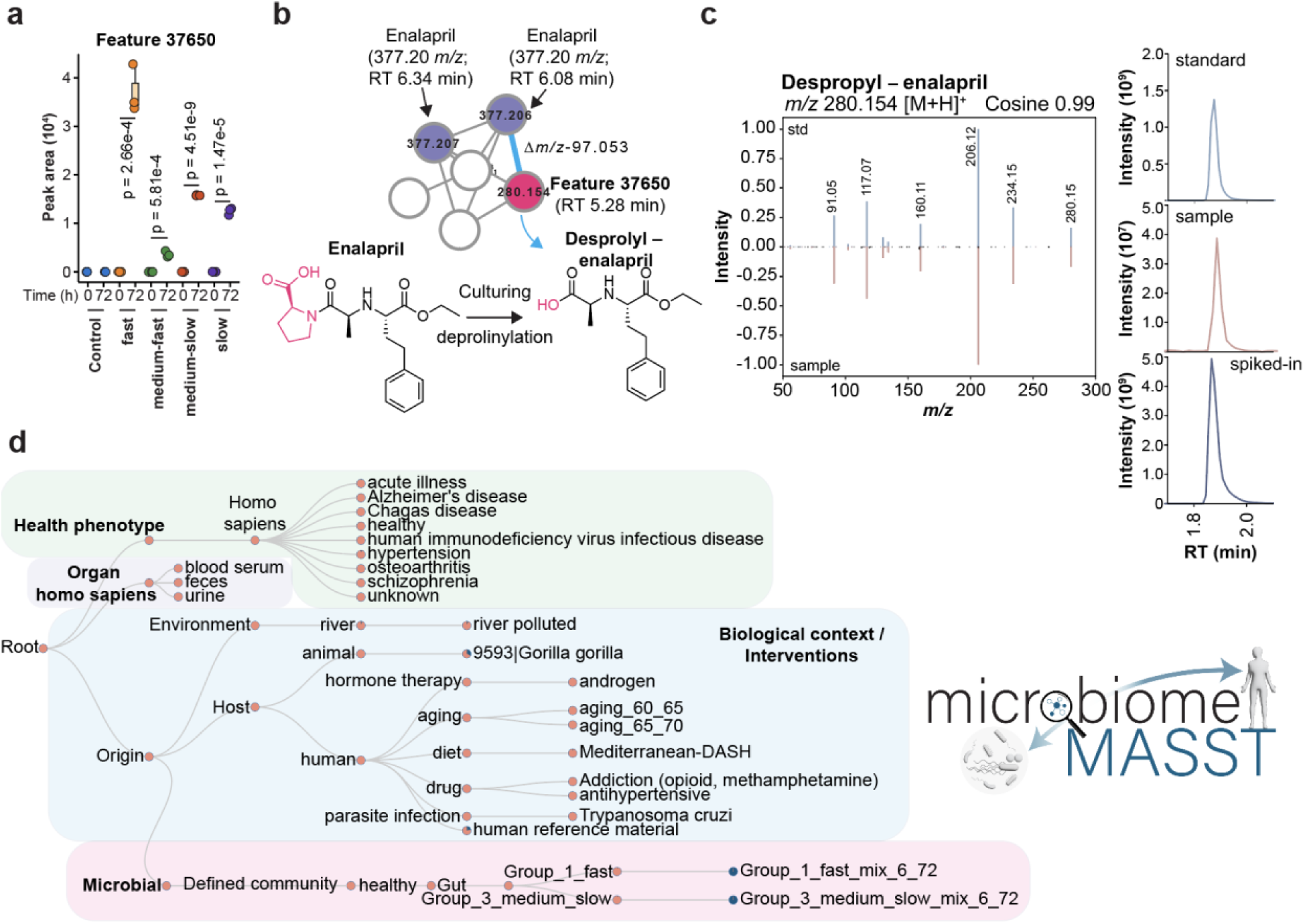
Discovery of a gut microbiota-derived metabolite of an angiotensin-converting enzyme inhibitor across human diseases. **a)** Boxplots showing the increase in the peak area for *m/z* 280.154 (RT 5.28 min) from baseline to 72 h across fast, medium-fast, medium-slow, and slow growers, compared to the medium control. **b)** Molecular network showing the feature with a *m/z* 280.154 (RT 5.28 min) clustering with enalapril, with a −97.053 Da delta mass consistent with enalapril deprolylation. **c)** Mirror plot showing MS/MS spectral similarity between the synthetic standard and the biological sample and extracted ion chromatogram from the standard versus the biological sample. **d)** MicrobiomeMASST results: the metabolite was detected in community culture of fast growers (Group1) and medium-slow growers (Group3); in aging and diet human cohorts, a human antihypertensive drug treatment dataset, zoo animals and polluted rivers, and was observed in feces, urine, and serum, and in several health phenotypes including osteoarthritis, Alzheimer’s disease, and IBD.

Using microbiomeMASST, we then tracked this metabolite across contexts: it was found in fast-growing bacteria (group 1; **Supplementary Table S2**) and medium-slow growers (group 3; **Supplementary Table S2**), in human antihypertensive cohorts, aging studies, in a polluted river, and unexpectedly in zoo gorillas (*Gorilla gorilla*). Conversations with zoo keepers revealed that enalapril is administered to gorillas^49^, and some zoos have specific usage guidelines due to potential side effects such as hepatoxicity^50^. Detection in feces, serum, and urine is consistent with systemic circulation and renal elimination, and the MS/MS to this metabolite was found across diverse health contexts such as osteoarthritis, IBD, and Alzheimer’s disease **(Fig. 3d)**.

Using microbiomeMASST, we were therefore able to translate this *in vitro* observation into human biology, without recruiting volunteers undergoing enalapril treatment or performing dedicated mouse experiments, efforts that are costly and time-consuming without guarantee of success. To further extend this finding across human-microbiome-drug systems, we reanalyzed a publicly available human aging cohort dataset ([MSV000097935](https://massive.ucsd.edu/ProteoSAFe/dataset.jsp?task=d6e24977a254438a88b0ec4a5f3adfe3)), in which desprolyl-enalaprilat was quantified in human serum at 0.85 µM, demonstrating the systemic circulation of this microbial metabolite in humans. Feature-based molecular networking highlighted a cluster containing desprolyl-enalapril adjacent to related angiotensin-converting enzyme inhibitors (ACE-inhibitors; trandolapril, quinapril) **(Fig. 4a)**. Annotation propagation further identified the active drug form - enalaprilat - in its desprolinylated form (Proline amide hydrolyzed-enalaprilat), consistent with either hydrolysis of the ethyl ester of the enalapril or direct desprolinylation of enalaprilat **(Fig. 4a; Supplementary Figure 5)**.

**Fig. 4.**
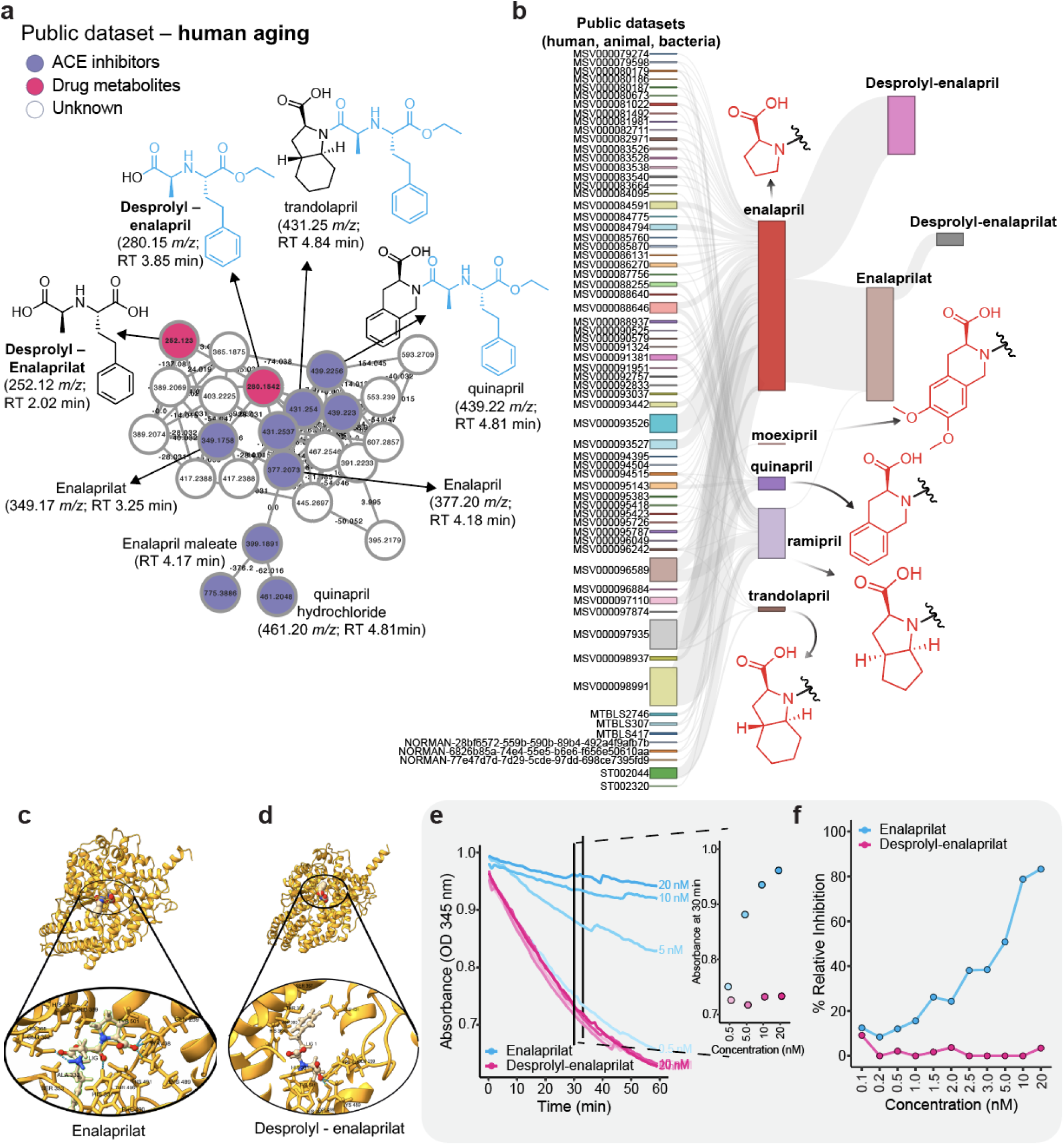
Gut microbes can inactivate the antihypertensive drug enalapril(at). **a)** Molecular network of the healthy human aging dataset [MSV000097935](https://massive.ucsd.edu/ProteoSAFe/dataset.jsp?task=d6e24977a254438a88b0ec4a5f3adfe3) shows associations of the drug metabolite to many angiotensin-converting enzyme (ACE) inhibitors. **b)** Sankey visualization showing the unique association of the desprolyl-enalapril to the prodrug enalapril and the association of desprolyl-enalaprilat with enalaprilat (active drug) compared with the other ACE inhibitors (moexipril, quinapril, ramipril, and trandolapril). **c)** Enalaprilat bound to angiotensin-converting enzyme 1 shows five hydrogen bonds (dotted blue lines) within 5Å involving Tyr498, Lys489, Gln259 (backbone), His331, and Glu362. The enlarged view is presented in **Supplementary Figure 6a,d**. Angiotensin-converting enzyme 1 (ACE1) bound to desprolyl-enalaprilat retains only two hydrogen bonds (Tyr498 and Lys489), indicating loss of anchoring interactions in the P2’ subsite. The enlarged view is presented in **Supplementary Figure 6a,b**. **e**) ACE1 enzymatic activity measured over time in the presence of increasing concentrations (0.5-20 nM) of enalaprilat (cyan) or desprolyl-enalaprilat (magenta). Vertical lines mark the 30-min time point used for comparison (right plot). Absorbance decreases with increasing enalaprilat concentration but remains unchanged with desprolyl-enalaprilat, indicating loss of inhibitory activity. **f)** Percentage of relative inhibition of ACE1 activity calculated from reaction slopes in the linear range (10-30 min). Enalaprilat shows a dose-dependent inhibition, whereas desprolyl-enalaprilat shows minimal inhibition across all tested concentrations.

Because several ACE inhibitors share a conserved scaffold with different side-chains, we next sought to determine whether this antihypertensive drug metabolite–or its active form–co-occurs only with enalapril(at) or also with other ACE inhibitors in human, bacterial, or animal datasets. To address this, we leveraged FASST to identify and compile all the datasets across four metabolomics repositories (GNPS/MassIVE^24^, MetaboLights^28^, Metabolomics Workbench^29^, and NORMAN/DSFP^51^) that contained matches to antihypertensive drugs sharing the same scaffold as enalapril(at), including despropyl-enalapril(at) (drug metabolite), moexipril, quinapril, ramipril, and trandolapril **(Fig. 4b)**. Co-occurrence network analysis revealed that the antihypertensive drug metabolites appear specifically with the parent drug enalapril and enalaprilat, potentially indicating specificity for cleavage of the proline moiety, but not when other chemical groups occupy this position (odds ratio [OR] = 91.04, 95% CI = 15.82–3576.66, Fisher’s exact p value = 1.77 × 10^−25^) **(Fig. 4b; Supplementary Figure 7)**. These repository-scale analyses support a microbiome-mediated deprolylation pathway detectable across multiple biological contexts and health conditions, showing how microbiomeMASST can translate *in vitro* observations to human datasets.

We next asked whether this microbiome-mediated drug modification of enalapril(at) compromises its ability to inhibit angiotensin-converting enzyme 1 (ACE1), given that the ester warhead essential for covalent binding remains intact. To structurally model this effect, we first built an *in silico* predictive protein-ligand model of ACE1 (PDB 6F9R) and co-folded enalaprilat (active drug) and desprolyl-enalaprilat using Boltz-2^52^ (**Fig. 4c-d**). In the enalaprilat co-folding model, five hydrogen bonds form within 5Å ligand oxygens to Tyr498-OH, Lys489-NH3+, and the Gln259 backbone, with additional contacts to His331 and Glu362. In contrast, loss of proline moiety leaves only two hydrogen bonds (Tyr498 and Lys489) (**Fig. 4c-d**). This reduction in the H-bonding network, along with removal of the proline residue, supports a model in which desprolyl-enalaprilat exhibits diminished ACE1 inhibitory activity. To provide support experimentally, we performed an ACE1 enzymatic inhibition assay by adding enalaprilat or desprolyl-enalaprilat to compete with the substrate for the active site of the ACE1 enzyme. While enalaprilat effectively inactivated ACE1 at all tested concentrations, desprolyl-enalaprilat showed no inhibitory effect (**Fig. 4e-f**).

## Discussion

MicrobiomeMASST provides a microbiome-focused, purpose-built analysis ecosystem designed to contextualize microbial metabolites across controlled experiments and complex biological systems. By enabling network-based linking of MS/MS spectra with experimentally validated microbial transformations and metadata spanning hosts, sample types, interventions, and culture conditions, this framework overcomes a central limitation of metabolomics studies that analyze datasets in isolation. Instead, microbiomeMASST allows microbial chemistry to be interpreted at repository scale, revealing recurring metabolic patterns, candidate producers, and biologically meaningful contexts that are otherwise difficult to resolve.

Applying this framework to bile acid metabolism uncovered a diverse and previously underappreciated landscape of microbially-conjugated bile acids, including conjugates with polyamines, diamines, decarboxylated amino acids, and dipeptides. MicrobiomeMASST connected previously reported bile acid–polyamine conjugates to specific bacterial producers, such as *Thomasclavelia spiroformis*, providing direct evidence that these molecules are synthesized by bacteria rather than arising solely from host or dietary processes. Beyond confirming known chemistry, culture-based bile acid supplementation experiments revealed previously unrecognized conjugates, including cholyl–ala–ala and cholyl–cystine, which were subsequently traced across bacterial cultures and digestive-tract–associated samples from humans and animals. The detection of these metabolites across systems highlights conserved, yet species- and location-specific, microbial bile acid transformations, for example, the restriction of cholyl–cystine to upper gastrointestinal samples suggests spatial organization of microbial bile acid conjugation.

Notably, the identification of bile acid–dipeptide conjugates raises the possibility that gut microbes may exploit bile acids as molecular carriers. Dipeptides are not actively transported across the intestinal epithelium, suggesting that bile acid conjugation could represent a microbial strategy to facilitate peptide transport beyond the gut. Within this context, *Intestinimonas butyriciproducens*, identified through microbiomeMASST as a producer of dipeptide bile acid conjugates, emerges as a candidate of interest for microbiota-based therapeutic strategies targeting metabolic disease^53–55^.

Within the same framework, microbiomeMASST also linked MS/MS evidence of drug metabolites across bacterial cultures and human samples (urine, blood, and feces), enabling the identification of microbial proline hydrolysis of the ACE prodrug enalapril. Enalapril is administered as an ester prodrug that is normally hydrolyzed to the active form enalaprilat. By integrating structure-based protein–ligand modeling and biochemical assays, these observations support a role for gut bacteria in drug inactivation, providing a mechanistic basis for interindividual variability in therapeutic response. Indeed, clinical studies comparing ACE inhibitors have reported differential antihypertensive efficacy between prodrugs. For example, in a randomized double-blind trial of patients with hypertension, lisinopril produced a greater and more sustained reduction in blood pressure than enalapril across a 12-week treatment period^56^. In heart-failure patients, switching from enalapril to the longer-acting ACE inhibitor perindopril was associated with significant improvements in functional status, blood pressure, and markers of left-ventricular remodeling, highlighting differences in ACE inhibitors^57^. Such differences are typically attributed to pharmacokinetic, dosing, or tissue penetrations; our findings indicate that gut microbial metabolism of enalapril may represent an additional, previously unrecognized layer of interindividual variability. In this framework, microbial hydrolysis or inactivation of enalapril prior to host conversion could limit systemic exposure to the active drug in a subset of individuals, contributing to heterogeneous clinical responses.

Together, these examples illustrate how microbiomeMASST enables hypothesis generation at the interface of microbial metabolism and host-relevant biology. The resulting predictions are directly actionable, guiding experimental follow-up and validation, as demonstrated in this work. In its current implementation (version 1), the microbiomeMASST network graph focuses on interventions and conditions with strong microbiome relevance, as well as synthetic microbiome systems for which well-curated untargeted metabolomics data and rich metadata are available, including gut, skin, mucosal, and plant-associated environments. Other microbiomes, such as rhizosphere and food-associated systems, are not yet comprehensively represented and will be incorporated in future expansions. Rather than functioning as a standalone discovery tool, this expandable framework integrates observational evidence across datasets, organisms, and experimental systems, enabling the prioritization of biologically and clinically relevant microbial chemistry for downstream mechanistic investigation.

### Future perspective and metagenomics data integration

Looking forward, microbiomeMASST provides the foundation to readily integrate metabolomics with metagenomics data. One promising direction is to link any MS/MS spectra from any bacterial culture with publicly available sequencing data, which could enable systematic association of microbial metabolites with putative producers across diverse ecosystems and their native complex systems, such as human samples. MicrobiomeMASST integration with resources such as redbiom^58^ and Qiita^59^ would allow metabolomics-first hypotheses to be evaluated in the context of vast genomic datasets (as of 11/2025, the public redbiom cache includes 318,955 samples from 561 studies for 16S V4 rRNA gene amplicon sequencing and 59,544 samples from 69 studies for whole-genome sequencing (WGS) with preparation and sample metadata), supporting multi-level inference. Within microbiomeMASST, each strain is associated with an operational genomic unit (OGU) corresponding to its Web of Life 2 (WoL2) identifier^48^. When the strain-level OGU is not available, the nearest taxon is assigned.

For example, we reported a previously unexplored microbially conjugated bile acid cholyl–ala-ala for which one producer is *E. faecalis* (OGU G000007785). Using redbiom^58^, this microbe can be traced within the same filenames, in which paired-omics data were generated and made publicly available on MassIVE and Qiita^49^, providing strong evidence that it is a plausible *in situ* producer and representing a critical step toward causal mechanistic follow-up studies.

More broadly, continued expansion of public metabolomics datasets and metadata deposition will further enhance the interpretability of microbial chemistry across hosts, environments, and clinical interventions. By enabling rapid cross-study contextualization, microbiomeMASST establishes a scalable framework for transforming dispersed metabolomics data into biological knowledge.

## Methods

### Microbiome-based network graph creation

A new microbiome-focused network graph has been built, improving the metadata capture system allowing standardized biological context and interventions of metabolome datasets. Public studies from metabolomics repositories GNPS/MassIVE, Metabolomics Workbench, and Metabolights were manually filtered and classified for microbiome research into categories currently not captured within current metadata ontologies like PanReDU. Categories includes genetic models (e.g., Alzheimer’s mouse model 3xTG and 5xFAD; diet interventions in human and mouse (e.g., low-fat/high-fat, high fibre, beef, and anti-inflammatory), sleep disruption, aging, germ-free, drug treatments. Health conditions and tissues/biofluids were manually curated and complemented using the PanReDU framework. Untargeted metabolomics data on microbial cultures were either acquired for this study (e.g., gut and skin microbes) or imported from the public domain (e.g., nasal, phyllosphere). Microbes were cultured individually or in community with and without building blocks to reveal a broader functional capacity which includes bile acids, 164 drugs and 268 food biomarkers and complex mixtures. In total, microbiomeMASST contains 467 datasets and 144,424 LC-MS unique files.

### Bacterial strains and culture conditions

#### Gut synthetic communities

Bacteria and community composition used throughout this study are listed in **Supplementary Table S2**. Bacteria cultures were started from glycerol stock at −80 ℃ and incubated in an anaerobic chamber (20% CO_2_, 5% H_2_, and 75% N_2_) in a modified BHI medium with pH adjusted to 7.5 with NaOH 5N **Supplementary Table S3**. One hundred and forty drugs were supplemented at a final concentration of 4 µM and drug details are provided in **Supplementary Table S4**. A second drug experiment supplemented at a final concentration of 2 µM was also conducted with 20 drugs with communities and details can be found in **Supplementary Table S4**. Details on food biomarker standards and complex food mixture are provided in **Supplementary Table S5**. Bacterial monocultures were normalized to an OD_600_ of 0.02, cultured into a 96-well plate with edges filled with sterile medium to prevent evaporation, and incubated for 48-72 h.

#### Crohn’s microbial community (cCOM)

The Crohn’s disease microbial community (cCOM) is composed of 12 keystone species isolated from patients with Crohn’s disease **Supplementary Table S2**. All strains were cultured individually and as 12-member community in brain heart infusion supplemented (BHIS) under anaerobic conditions (10% CO_2_, 7.5% H_2_, and 82.5% N_2_), except for *Enterocloster bolteae* and *Bacteroides caccae*, which were grown in Reinforced Clostridial Medium (RCM) and Brucella with pyruvate, taurine, and iron (BruPT-Fe), respectively (**Supplementary Table S7**). Bacteria cultures were incubated for 72 h with and without 5-aminosalicylic acid (5-ASA; 200 µM), 3-phenylpropionic acid (200 µM), and four taurine bile acid conjugates (TCA, TLCA, TDCA, TCDCA), each at 50 µM. Bacteria were also extracted at time 0 h to establish a baseline.

#### Metabolite extraction

After incubation, bacterial cells were extracted with a ratio sample to solvent of 1:4 with either 50% or 80% of pre-chilled MeOH/H_2_O and incubated overnight at 4 ℃. Samples were centrifuged at 2,500 rpm for 10 min and the supernatants were transferred to a new plate before being dry in a CentriVap overnight. Samples were stored at −80 ℃ until LC-MS/MS analysis.

#### LC-MS/MS data acquisition

Samples were resuspended in 80% pre-chilled MeOH/H_2_O containing sulfadimethoxine as internal standard and incubated at −20 ℃ overnight before being centrifuged either at 2,000 rpm for 20 min or 21,130 x g for 10 min depending if the samples were prepared in plates or in vials, respectively. Before injection, samples were randomized and two different mass spectrometers were used to acquire data. A Q-Exactive Orbitrap and an Orbitrap Exploris 240, both coupled to a Vanquish UHPLC system (Thermo Fisher Scientific). A 5 µL was injected into the LC system and a polar C18 (Kinetex, 100 x 2.1 mm, particle size 2.6 µm, pore size of 100 Å, Phenomenex) was used for the chromatographic separation. The mobile phase consisted of solvent A (H_2_O + 0.1% FA) and solvent B (ACN + 0.1% FA). The samples were placed in a pre-cooled chamber (4 ℃), injected in a heated column at 40℃, and eluted using a flow rate of 0.5 mL/min using the following gradient elution program: 0.5-1 min, 5 %B; 1-7.5 min, 5-40 %B; 7.5-8.5 min, 40-99%B; 8.5-9.5 min, 99 %B; 9.5-10 min, 95-5 %B; 10-10.5 min, 5 %B; 10.5-10.75 min, 5-99 %B; 10.75-11.5 min, 99 %B; and re-equilibration step from 11.5-12 min, 99-5 %B.

For data acquired on Q Exactive Orbitrap, data-dependent acquisition (DDA) of MS/MS spectra was performed in positive ionization mode. ESI parameters were set to 53 L/min, auxiliary gas flow rate 14 L/min, sweep gas flow rate 3 L/min, spray voltage 3.5 kV, inlet capillary set to 269 ℃, and auxiliary gas heater set to 438 ℃. The mass scan range was set to 100-1000 *m/z* with a resolution of 35,000 at *m/z* 200. The automatic gain control (AGC) target was set to 1E6, with a maximum injection time set to 100 ms. Spectral data was acquired in profile mode. Up to 5 MS/MS spectra per MS1 scan were acquired with a resolution of 17,500 at *m/z* 200 of with one microscan. AGC target was set to 5E5, with a maximum injection time of 150 ms. The isolation window was set to 1 *m/z* with isolation offset set to 0 *m/z*. The normalized collision energy was acquired with a stepwise increase of 25, 40, and 60%. The apex trigger was set to 2 - 15 s with a dynamic exclusion of 5 s, and exclude isotope set to ON.

For data acquired on Orbitrap Exploris 240, DDA method was performed in positive ionization using the following parameters: 3.5 kV spray voltage, Sheath Gas flow rate 50 Arb, auxiliary Gas 10 Arb, sweep Gas 1 Arb, ion transfer tube temperature set to 325 ℃, and Vaporizer temperature set to 350 ℃. The mass scan range was set to 150-1500 *m/z* with a resolution of 60,000 at *m/z* 200 with one microscan. Rf Lens was set to 70%, AGC was set to Standard, and the data acquired in profile mode. An intensity of 1E4 was set to trigger a DDA scan. Dynamic exclusion was set to custom with the following parameters: precursor ion was excluded after 2 times if it occurs within 3 s for a duration of 4 s. The isolation window was set to 1 *m/z* with the isolation offset set to OFF. Normalized collision energy was set to a stepwise increase of 25, 40, and 60%. Top 10 MS/MS spectra per MS1 scan were acquired with a resolution of 22,500 at *m/z* 200. Scan range was set to auto with AGC target set to custom (200%), and maximum injection time set to auto with one microscan. For data acquired using the AcquireX Deep scan mode, two blanks were injected with the second one being used as a reference and four pool injections were performed to create an exclusion list.

#### Quantitation and retention time matching

Quantitation of desprolyl-enalaprilat was performed from a serum sample of an aging cohort (UCSD IRB 960019; UCSD IRB 040165; UCSD IRB 130757). Scripts used for quantitation are available in the GitHub repository (see data availability section). The LC-MS/MS method used for quantitation is the same as described above. The analytical method followed the International Conference on Harmonization (ICH) guidelines^61^. The method was validated using the following parameters: specificity, limit of detection (LOD), limit of quantitation (LOQ), accuracy, linearity, and precision. Briefly, a stock solution of 1 mM of desprolyl-enalaprilat was prepared in 50% MeOH/H_2_O, followed by a serial dilution to get final concentration range of 0.05 μM to 4 μM for the calibration standards (calibrators). Each calibrator was injected in triplicate. Peak area for desprolyl-enalaprilat and sulfadimethoxine (internal standard) were extracted using Skyline^62^. A polynomial equation was obtained and the correlation coefficient was calculated for desprolyl-enalaprilat to be 0.996. The limit of detection (LOD) was determined using by the mean of the slopes (S) and the standard deviation of the y intercept, lower limits of quantification (LLOQ) is 3.3* LOD. The accuracy and precision of the calibration curve was validated by spiking serum into 100 nM and 4 μM desprolyl-enalaprilat to account for matrix effect, for intra-day, n = 6, the accuracy and precision was 90.5% and CV% 5.5 and inter-day, n = 3, 90.7% and CV% 4.6. Peak area of desprolyl-enalaprilat at each concentration was normalized using sulfadimethoxine. Serum sample from humans was retrieved from an aging cohort and injected (n = 3) using the validated method. The absolute concentration was calculated to be 0.85 µM considering the factor of dilution used during the extraction process (dilution 0.5x). The specificity was assessed by blank injection with internal standard sulfadimethoxine and injection of the desprolyl-enalaprilat (n = 3) followed by the calculation of the relative standard deviation (RSD) based on the biological sample. The retention time of desprolyl-enalaprilat in both the biological sample and the authentic standard was 1.74 min. Raw data for the quantitation can be found in **Supplementary Table S9**. Retention time matching and MS/MS spectral similarities were performed between the biological samples (e.g., bacterial cultures, human urine samples) and the crude synthetic standard or combinatorial synthesis available on MassIVE ([MSV000100461](https://massive.ucsd.edu/ProteoSAFe/dataset.jsp?task=84923b9049784897af321b0764a203e9)).

#### Untargeted metabolomics data processing

The raw files generated by the mass spectrometers were converted in an open-access format mzML using MSConvert^63^ (version 3.0.25042 64-bit) and datasets acquired in this study were uploaded to MassIVE (see data availability section). Feature detection and extraction were performed in MZmine 4 using the batch processing mode. An example of the batch processing file (.mzbatch) can be found in the associated GitHub repository. Briefly, the data was imported using MS1 and MS/MS mass detection modules using the factor of the lowest final and noise factor set to 4 and 2, respectively. The chromatogram builder was used with a minimum consecutive scans of 4 with a minimum intensity of 3E4, a minimum absolute height of 1.5E5, and a *m/z* tolerance of 10 ppm. Smoothing was applied using Savitzky Golay algorithm, before using the local minimum feature resolver module which had the following parameters: chromatographic threshold of 85%, minimum search range 0.05 min, an absolute height of 1.5E5, min ratio peak top/edge of 2, and a minimum scans of 4. Isotope (13C) was filtered followed by isotope finder. Features were aligned using a 10 ppm *m/z* tolerance with weight *m/z* of 3, and a retention time of 0.3 min. Features detected in fewer than two samples and less than 10% of all samples were filtered out, unless they triggered an MS/MS event, before applying feature finder. MetaCorrelate module was added with a retention time tolerance of 0.06 min with feature height correlation set to minimum of samples 2 and a minimum Pearson correlation of 70%. Ion identify molecular networking was performed before exporting the tables. GNPS and SIRIUS were used to export the feature table and the mgfs, necessary for downstream analyses. For quality control assessment, the legacy MZmine 2 module was exported as well.

#### Untargeted metabolomics data analysis

The feature tables were imported into RStudio (version 4.5.1) for downstream analysis. Data quality was evaluated by analyzing the retention times and the peak areas of the internal standard (IS) across all samples and by computing the coefficient of variance of the IS in the biological sample and across a mixture of six standards (amytriptiline, sulfamethizole, sulfamethoxine, sulfadimethoxine, coumarin 314, and sulfachlopyridazine) present in a Qcmix sample injected every 10 samples. Blank subtraction was performed by removing features in which their peak areas were not at least five times higher than the poolQc and Qcmix samples. To be displayed in the radial network plots, the peak area of a feature needs to be at least three times higher than in the control condition or at least increasing three times from baseline (0 h) to 72 h. Features matching water-loss delta masses were filtered out.

#### Analysis of public dataset from GNPS/MassIVE

All datasets analyzed in this study were imported from GNPS/MassiVE and processed using MZmine 4^64^. The output from the batch files generates several output files such as quality control (module legacy), feature table (quant.csv), and a mgf file containing the spectra. The quantification table and the spectra mgf file are both imported into GNPS2 to perform feature-based molecular networking (FBMN) workflow. The parameters used for all FBMN jobs are precursor ion tolerance and fragment ion tolerance of 0.02 and filters set to OFF. The networking parameters and library search parameters vary; cosine 0.6 or 0.7 and numbers of matched peaks between 4 and 6. The settings have been chosen based on the chemical classes and mass spectrometers used to collect data currently available in the public domain.

Culturing of gut microbes with bile acids and polyamines ([MSV000095648](https://massive.ucsd.edu/ProteoSAFe/dataset.jsp?task=a2b196ed4b0a4236ba7b0fdddbd0ebc7)): https://gnps2.org/status?task=a6c94de887d841a199860d8e67a531a6

Eight gut microbes in monoculture with bile acids and drugs ([MSV000097269](https://massive.ucsd.edu/ProteoSAFe/dataset.jsp?task=a0669877adb44e2d8681d8e2cdfc5720)): https://gnps2.org/status?task=ec12e706a21d4d178983c9380174fb82

140 drugs with communities ([MSV000096589](https://massive.ucsd.edu/ProteoSAFe/dataset.jsp?task=1ab5cea1a134430a9b0899fe50a278aa)): https://gnps2.org/status?task=bdd2bea069514fa7871cfe517c388df3

Aging study ([MSV000097935](https://massive.ucsd.edu/ProteoSAFe/dataset.jsp?task=d6e24977a254438a88b0ec4a5f3adfe3)) https://gnps2.org/status?task=d51a47b836024d518886ec3d94463d01

### Analysis of public data using FASST

FASST, a fast version of MASST, was used to retrieve all the spectra found in the public domain. The list of USIs matching the antihypertensive drugs is provided in Supplementary Table S6. The FASST batch workflow was run with the python script available at https://github.com/robinschmid/microbe_masst. The library “metabolomicspanredu_index_nightly” was used to query the list of USIs, with a precursor and fragment ion tolerance of 0.01, minimum cosine similarity of 0.7, and a minimum of matched peaks of 3. The software R studio (version 4.5.0) was used for downstream analysis. Briefly, FASST search results were merged into a single dataframe in RStudio. Matches to non-human, bacteria, and animals were filtered out from the analysis based on manual inspection of all datasets retrieved from the FASST search. A co-occurrence analysis of both the parent drugs and the drug metabolites across files was performed.

### *In silico* protein-ligand co-folding assay

The co-folded structures of enalaprilat, desprolyl-enalaprilat, and angiotensin-converting enzyme 1 (ACE1) were predicted using the Boltz-2 foundational model^52^ implemented on the Rowan Scientific platform (https://www.rowansci.com; accessed 10/12/2025). For each inference, the SMILES representations of enalaprilat and desprolyl-enalaprilat were provided together with the amino acid sequence of ACE, using the default Boltz-2^52^ hyperparameters. The resulting co-folded complexes were analyzed in ChimeraX (version 1.10rc202505280131)^65^ to identify ligand–protein hydrogen bonds using the default geometric criteria and to quantify ligand–protein contacts within a 5 Å cutoff. These analyses were subsequently used to generate visualizations of the ligand–binding site.

### Angiotensin-converting enzyme 1 activity assay

ACE1 inhibitory activity was measured using the colorimetric ACE1 assay kit (Abcam; ab283407). Test compounds were dissolved in the ACE assay buffer (1 µM) and serially diluted to final concentration ranging from 0.1 nM to 20 nM. Each compound was incubated with the enzyme at 37 ℃ for 15-20 min in the 96-well plate, followed by the addition of the reaction mix according to the manufacturer’s protocol. The absorbance of each well was immediately measured at an optical density (OD) of 345 nm for 60 min using a plate reader. Absorbance data and corresponding time values (ms) were exported and analyzed in RStudio. Briefly, the slope was calculated for each well in the linear range (10-30 min). The enzyme control (EC) wells were used to define the baseline and the mean was used to compute the relative inhibition for each sample constrained between 0 and 100%, again according to the manufacturer’s protocol. For kinetic representative traces (0.5 nM - 20 nM), raw absorbance was plotted with the 30 min time marked.

## Data availability

All data within microbiomeMASST are publicly available and accessible in the three main metabolomics repositories (GNPS/MassIVE, Metabolights, and Metabolomics Workbench). A list of all the datasets embedded in this resource is provided in **Supplementary Table S1**. Datasets generated in this study have been deposited in GNPS/MassIVE: [MSV000096589](https://massive.ucsd.edu/ProteoSAFe/dataset.jsp?task=1ab5cea1a134430a9b0899fe50a278aa) (140 drugs with communities), [MSV000098638](https://massive.ucsd.edu/ProteoSAFe/dataset.jsp?task=37fc7ed54e344f3290d154e85ba8a0b4) (dietary substances and drugs), [MSV000097877](https://massive.ucsd.edu/ProteoSAFe/dataset.jsp?task=8ff0b9d6e1d24bd388532f7922b59784) (SkinCom cultures), [MSV000097269](https://massive.ucsd.edu/ProteoSAFe/dataset.jsp?task=a0669877adb44e2d8681d8e2cdfc5720) (monocultures with drugs (3)), [MSV000097155](https://massive.ucsd.edu/ProteoSAFe/dataset.jsp?task=3c6c4a62651d4459bfc7f88c115fed1d) (microbes with bile acids), [MSV000095648](https://massive.ucsd.edu/ProteoSAFe/dataset.jsp?task=a2b196ed4b0a4236ba7b0fdddbd0ebc7) (monocultures with bile acids), and [MSV000095331](https://massive.ucsd.edu/ProteoSAFe/dataset.jsp?task=4ec7dab059204b1ea69613cbd8f56ce3) (20 drugs with communities). MicrobiomeMASST interface is accessible using the following link: https://masst.gnps2.org/microbiomemasst/. Representative videos are presented to help guide the user navigate microbiomeMASST.

Video 1 (N-acetyl spermidine conjugate): https://youtu.be/6KFjCKO_IWc

Video 2 (2,3-diaminopropionic acid conjugate): https://youtu.be/XcwbCSKClYM

Video 3 (Cholyl-ala-ala conjugate): https://youtu.be/xkZ8DVCT3NU

Video 4 (Cholyl-cystine conjugate): https://youtu.be/Ybaonh5b7Qo

Video 5 (deoxycholyl-phenylethylamine): https://youtu.be/G42IMBvw3Dk

## Code availability

All the scripts used to perform data analysis and generate figures are publicly available at https://github.com/VCLamoureux/microbiomeMASST.

## Acknowledgements

P.C.D appreciates the support by BBSRC/NSF award 2152526, National Institute of Health Sciences U24DK133658, R01DK136117 and Chan Zuckerberg Initiative. S.L. is supported by Research Council of Finland funding grant no. 363417. V.C.L. is supported by Fonds de recherche du Québec - Santé (FRQS) postdoctoral fellowship (335368) and from Natural Sciences and Engineering Research Council of Canada (NSERC) postdoctoral fellowship (598938). P.C.D and A.M.C-R. were supported by the Gordon and Betty Moore Foundation, GBMF12120 (https://doi.org/10.37807/GBMF12120). H.N.Z. was supported by the National Institute Of Environmental Health Sciences of the National Institutes of Health under Award Number K99ES037746. L.P. is supported by the University of California San Diego Medical Scientist Training Program (NIH/NIGMS T32GM154642). E.R.R. was supported by PSU/NIDDK (T32DK120509). A.D.P. was supported by the National Institute of Environmental Health Sciences (R35ES035027) and the USDA National Institute of Food and Federal Appropriations under Project PEN047702 and Accession number 7006412. A.T.A. was supported by the National Institute of Health (R35GM155026). S.F.H.S. is supported by the National Institute of Health (F32DE035388-01). S.G. is supported by a Medical University of South Carolina (MUSC) Odyssey Fellowship. M.A.E. is supported by the National Institute of Health (P30DK123704, P20GM120457, R35GM155451). The Texas Children’s Hospital Department of Pathology provides salary support to the bioanalytical chemistry staff members who work in The Virginia and L.E. Simmons Family Foundation Mass Spectrometry laboratory in the Texas Children’s Research Institute – Microbiome Center (T.D.H.). T.D.H. is supported by an Office of Research Infrastructure Programs (ORIP)-based S10 Award (NIH S10OD036416), the Texas Medical Center - Digestive Disease Center (NIH P30DK056338), and receives salary support from a P01 Program Project Grant (NIH P01AI152999). A.L.L. was supported by generous lab start-up funds from UC San Diego. A.E.S was supported by the Molecular Biophysics Training Grant, NIH Grant T32 GM139795. R.P.G. was supported by the Xunta de Galicia through the Postdoctoral Fellowship Program (IN606B-2024.22). AJAM and JB was funded by The Wellcome Leap Dynamic resilience program (co-funded by Temasek Trust).

## Author contributions

VCL and PCD conceptualized the project. VCL, CW, VD, KK, SZ, VG, AM, LC, ME, SR, and HG performed culturing experiments. VCL and JA acquired the LC-MS/MS data. VCL, SZ, SX, PK, YEA, and MW developed the computational pipeline and interface. AP, ZH, MP and SG, did the synthesis part of the project supervised by DS. VCL, SZ, RAM, and AJ did the *in silico* modeling prediction, IM and HG did the protein activity assay. VCL, HNZ, GA, AMCR, KEK, RPG, MSC, NS, JZ, ATA, LL, SD, SL, ERR, VN, AES, ALL, SFHS, SG, MAE, TDH, MM, SKM, HY, MG, ES, NHP, PS, EK, AKPS, DP, MR, ADP, SD, KZ, AJAM, JB, contributed data. All authors tested the microbiomeMASST interface. LP, YW, AG, DM, and RK developed redbiom and conducted analyses. VCL and PCD wrote the manuscript and all authors reviewed and approved the manuscript.

