## Supporting information for "A searchable metadata network graph for microbiome metabolomics"

**Disclosures**

P.C.D. is an advisor and holds equity in Cybele, BileOmix, Sirenas and a scientific co-founder, advisor, holds equity and/or received income from Ometa, Enveda, and Arome with prior approval by UC San Diego. P.C.D. also consulted for DSM animal health in 2023. R.K is a scientific advisory board member, and consultant for BiomeSense, Inc., has equity and receives income. He is a scientific advisory board member and has equity in GenCirq. He has equity in and acts as a consultant for Cybele. He is a co-founder of Biota, Inc., and has equity. He is a co-founder of Micronoma and has equity and is a scientific advisory board member. He is a board member of Microbiota Vault, Inc. He is a board member of N=1 IBS advisory board and receives income. He is a Senior Visiting Fellow of HKUST Jockey Club Institute for Advanced Study. The terms of these arrangements have been reviewed and approved by the University of California, San Diego in accordance with its conflict of interest policies. M.W. is a co-founder of Ometa Labs LLC. D.M. is a consultant for and has equity in BiomeSense, Inc. The terms of these arrangements have been reviewed and approved by the University of California, San Diego, in accordance with its conflict-of-interest policies. TDH is a member of the Editorial Advisory Board and is contracted as an Associate Academic Editor for a Cell Press journal called STAR Protocols. TDH is also a participant in the SCIEX Global Thought Leaders in Mass Spectrometry Program. L.L. is a cofounder and equity holder of NiMo Therapeutics, and holds equity of Mirvie. SKM is a co-founder of Vertero Therapeutics and Nuanced Health and declares no competing interests.

**Author contributions**

VCL and PCD conceptualized the project. VCL, CW, VD, KK, SZ, VG, AM, LC, ME, SR, and HG performed culturing experiments. VCL and JA acquired the LC-MS/MS data. VCL, SZ, SX, PK, YEA, and MW developed the computational pipeline and interface. AP, ZH, MP and SG, did the synthesis part of the project supervised by DS. VCL, SZ, RAM, and AJ did the *in silico* modeling prediction, IM and HG did the protein activity assay. VCL, HNZ, GA, AMCR, KEK, RPG, MSC, NS, JZ, ATA, LL, SD, SL, ERR, VN, AES, ALL, SFHS, SG, MAE, TDH, MM, SKM, HY, MG, ES, NHP, PS, EK, AKPS, DP, MR, ADP, SD, KZ, AJAM, JB, contributed data. All authors tested the microbiomeMASST interface. LP, YW, AG, DM, and RK developed redbiom and conducted analyses. VCL and PCD wrote the manuscript and all authors reviewed and approved the manuscript.

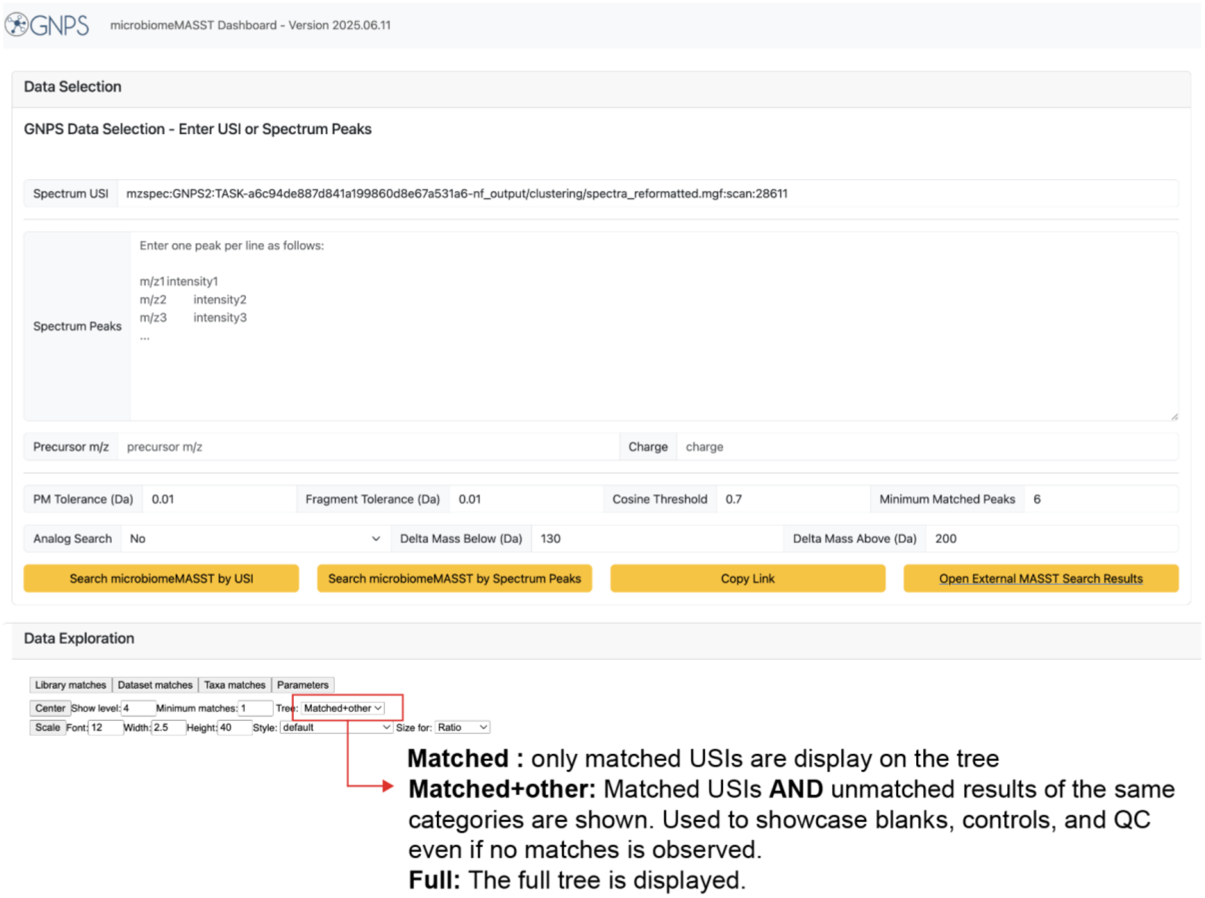


**Supplementary Figure 1 | MicrobiomeMASST user interface and advanced tree visualization options.** microbiomeMASST web app interface is accessible here https://masst.gnps2.org/microbiomemasst/. Users have two options to search data embedded in microbiomeMASST; entry of the Universal Spectrum Identifier (USI) or manual input of the spectrum. The latter requires manual entry of the precursor *m/z* and the charge. Search parameters are customizable by the users and includes precursor mass tolerance, fragment tolerance, cosine threshold, and number of matched peaks. After the search has been launched, under the “Data Exploration section”, users can explore the content and three tree visualization options are available. matched, matched+other, and full.

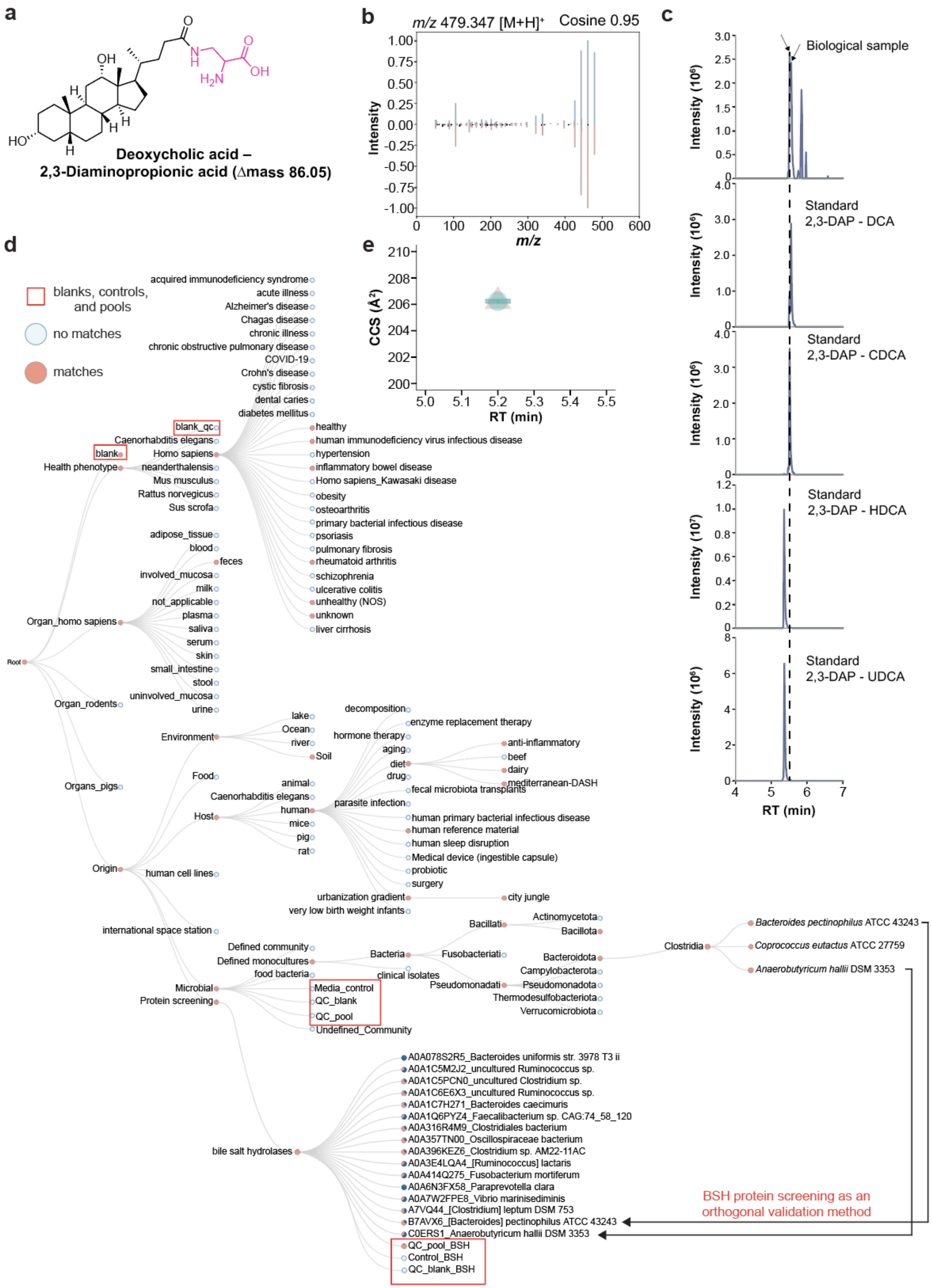
a
Deoxycholic acid –
2,3-Diaminopropionic acid (Δmass 86.05)
b
m/z 479.347 [M+H]+
Cosine 0.95
Intensity
m/z
c
Biological sample
Standard
2,3-DAP - DCA
Standard
2,3-DAP - CDCA
Standard
2,3-DAP - HDCA
Standard
2,3-DAP - UDCA
Intensity (10⁶)
Intensity (10⁷)
RT (min)
d
blanks, controls, and pools
no matches
matches
e
CCS (Å²)
RT (min)
Root
Health phenotype
blank
blank_qc
Caenorhabditis elegans
Homo sapiens
neanderthalensis
Mus musculus
Rattus norvegicus
Sus scrofa
acquired immunodeficiency syndrome
acute illness
Alzheimer's disease
Chagas disease
chronic illness
chronic obstructive pulmonary disease
COVID-19
Crohn's disease
cystic fibrosis
dental caries
diabetes mellitus
healthy
human immunodeficiency virus infectious disease
hypertension
inflammatory bowel disease
Homo sapiens_Kawasaki disease
obesity
osteoarthritis
primary bacterial infectious disease
psoriasis
pulmonary fibrosis
rheumatoid arthritis
schizophrenia
ulcerative colitis
unhealthy (NOS)
unknown
liver cirrhosis
Organ_homo sapiens
adipose_tissue
blood
feces
involved_mucosa
milk
not_applicable
plasma
saliva
serum
skin
small_intestine
stool
uninvolved_mucosa
urine
Organ_rodents
Organs_pigs
Origin
Environment
lake
Ocean
river
Soil
Food
Host
animal
Caenorhabditis elegans
human
mice
pig
rat
decomposition
enzyme replacement therapy
hormone therapy
aging
diet
drug
anti-inflammatory
beef
dairy
mediterranean-DASH
fecal microbiota transplants
parasite infection
human primary bacterial infectious disease
human reference material
human sleep disruption
Medical device (ingestible capsule)
probiotic
surgery
urbanization gradient
city jungle
very low birth weight infants
human cell lines
international space station
Microbial
Defined community
Defined monocultures
food bacteria
Media_control
QC_blank
QC_pool
Undefined_Community
Bacteria
Bacillati
Actinomycetota
Bacillota
Fusobacteriati
Bacteroidota
Campylobacterota
clinical isolates
Pseudomonadati
Pseudomonadota
Thermodesulfobacteriota
Verrucomicrobiota
Clostridia
Bacteroides pectinophilus ATCC 43243
Coprococcus eutactus ATCC 27759
Anaerobutyricum hallii DSM 3353
Protein screening
bile salt hydrolases
A0A078S2R5_Bacteroides uniformis str. 3978 T3 ii
A0A1C5M2J2_uncultured Ruminococcus sp.
A0A1C5PCN0_uncultured Clostridium sp.
A0A1C6E6X3_uncultured Ruminococcus sp.
A0A1C7H271_Bacteroides caecimuris
A0A1Q6PYZ4_Faecalibacterium sp. CAG:74_58_120
A0A316R4M9_Clostridiales bacterium
A0A357TN00_Oscillospiraceae bacterium
A0A396KEZ6_Clostridium sp. AM22-11AC
A0A3E4LQA4_[Ruminococcus] lactaris
A0A414Q275_Fusobacterium mortiferum
A0A6N3FX58_Paraprevotella clara
A0A7W2FPE8_Vibrio marinisediminis
A7VQ44_[Clostridium] leptum DSM 753
B7AVX6_[Bacteroides] pectinophilus ATCC 43243
C0ERS1_Anaerobutyricum hallii DSM 3353
QC_pool_BSH
Control_BSH
QC_blank_BSH
BSH protein screening as an orthogonal validation method

**Supplementary Figure 2 | MicrobiomeMASST network graph provides an orthogonal method confirming the microbial origin of deoxycholyl–2,3-diaminopropionic acid. a)** We discovered a novel microbially-conjugated bile acid deoxycholyl–2,3-diaminopropionic acid conjugate confirmed via **b)** MS/MS spectral similarities and **c)** retention time matching of authentic synthetic standard and biological sample. **d)** The tree visualization option "matched+other" was chosen to biologically contextualize deoxycholyl–2,3-diaminopropionic acid and demonstrate its absence in culture controls and blanks. Finally, an orthogonal protein screening assay of 126 bile salt hydrolases is embedded within microbiomeMASST for additional confirmation of microbially-derived metabolites. Arrows indicate co-eluting peaks. **e)** Experimental validation of 2,3-diaminopropionic acid using retention time and collision cross section (CCS) measurements confirming the annotation. The triangle represents the biological samples and the circle denotes the authentic standard. The error bar represents the standard deviation of three replicates.

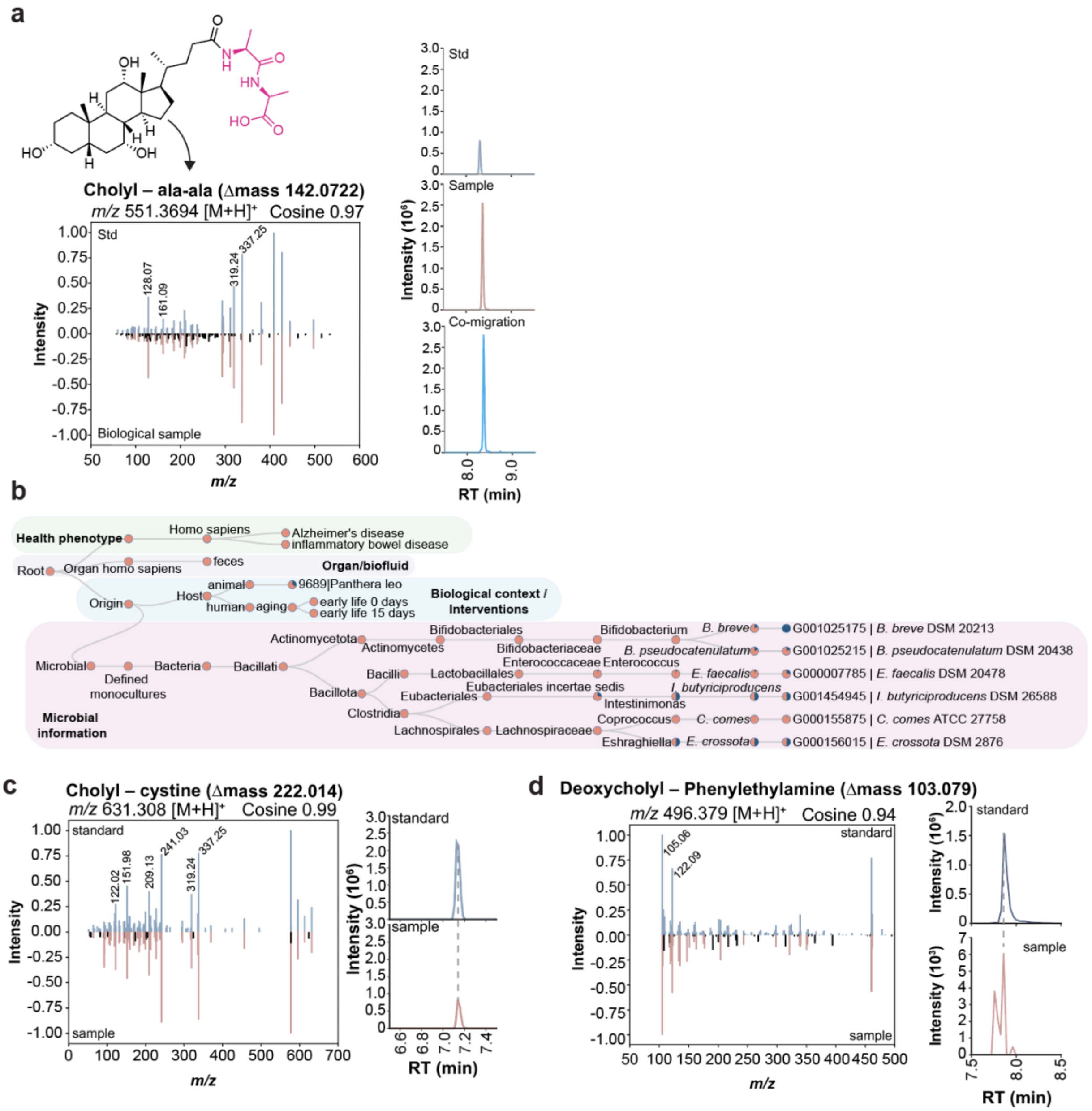


**Supplementary Figure 3 | Discovery and contextualization of cholyl - ala-ala. a)** Chemical structure of cholyl–ala-ala and associated mirror plot showing MS/MS spectral similarity and the extracted ion chromatogram from the standard versus the biological sample. **b)** MicrobiomeMASST results showing that cholyl–ala-ala was detected in six bacterial strains, in early-life, in an animal, in feces, across health phenotypes including Alzheimer's disease and IBD. **c)** MS/MS spectral matching and retention time alignment between the standard and the biological sample for cholyl–cystine and **d,** for deoxycholyl–phenylethylamine.

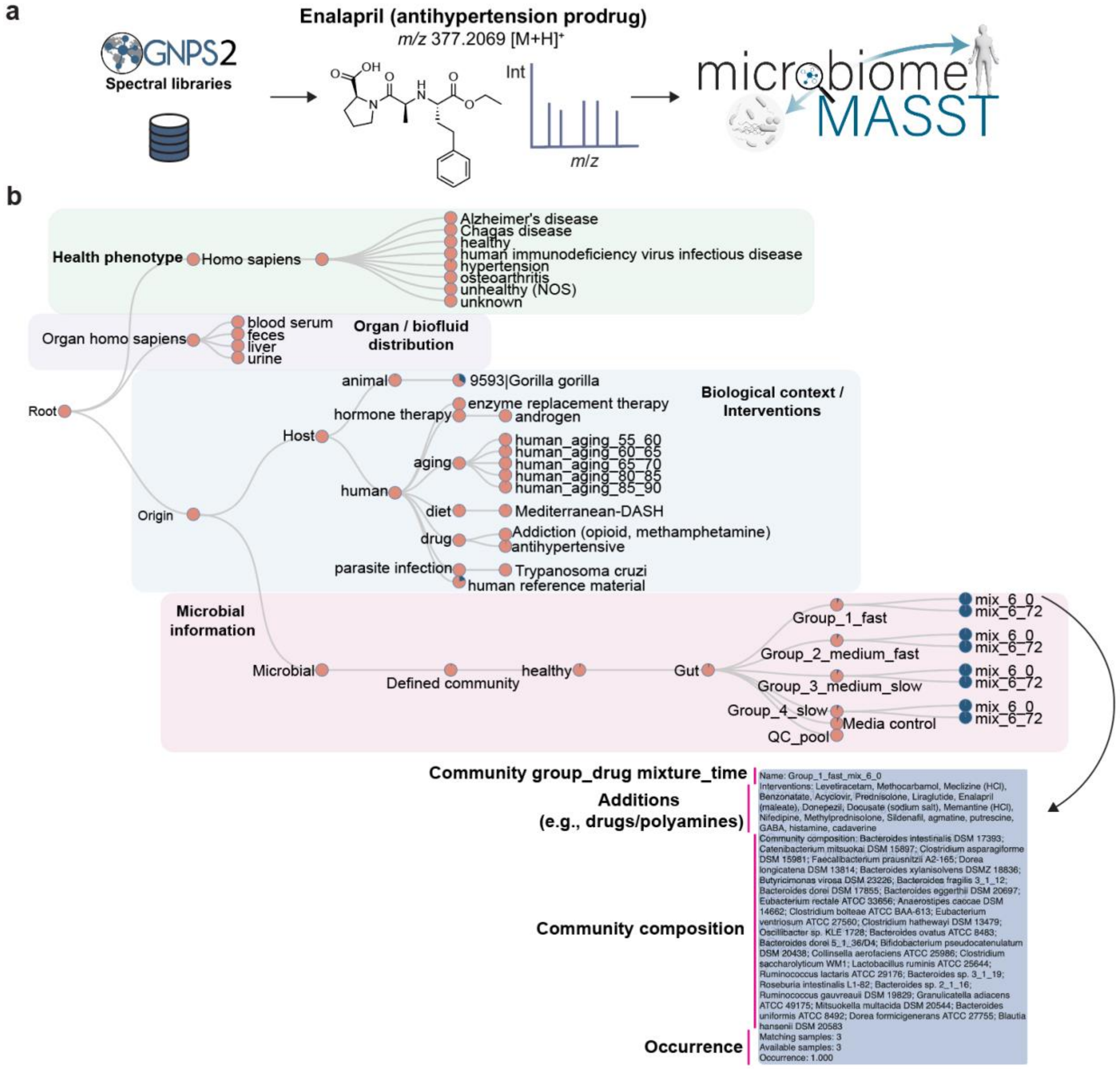


**Supplementary Figure 4 | Example to showcase how to interpret the data in the host-drug-microbe axis. a)** The universal spectrum identifier (USI) for enalapril was retrieved from GNPS2 spectral libraries and used as input in the microbiomeMASST web interface to query. **b)** MicrobiomeMASST results shown as a network graph of MS/MS matches across studies. Hovering over the nodes reveals information rich metadata including community group, drug mixture identifier, microbial incubation time (e.g., 0 h, 72 h). Moreover, it reports the intervention type (e.g., list of drugs, polyamines, bile acids) that was supplemented in the culture medium, the members of the community, and shows the occurrence.

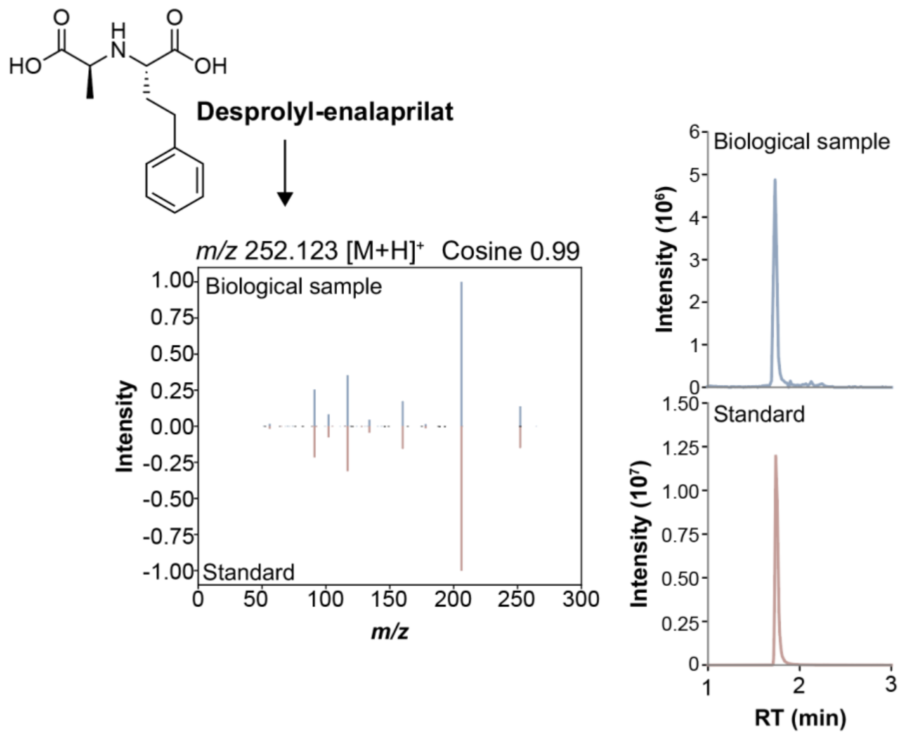


**Supplementary Figure 5 | Validation of desprolyl-enalaprilat in human serum sample.** Chemical structure of desprolyl-enalaprilat and associated mirror plot showing MS/MS spectral similarity and the retention time alignment between the biological sample and the authentic standard.

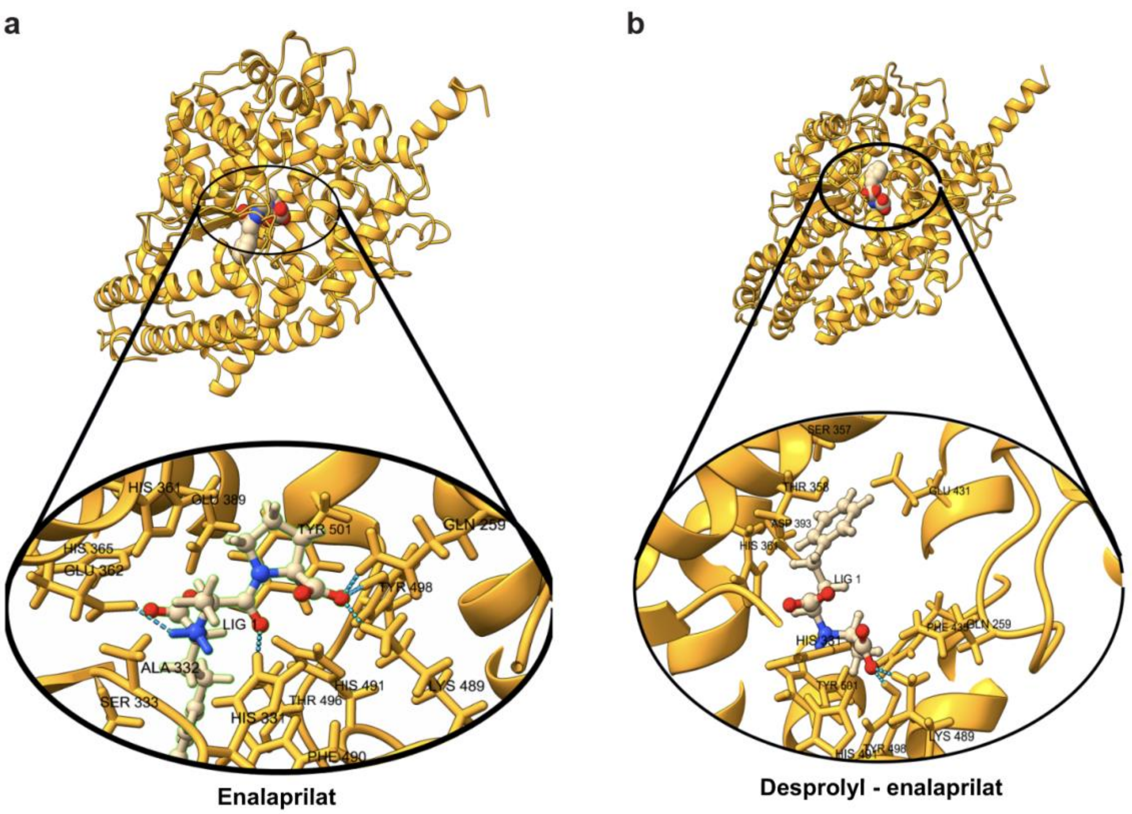


**Supplementary Figure 6 | *In silico* co-folding of ACE1 with enalaprilat and desprolyl-enalaprilat. a)** Enlarge view for enalaprilat and **b)** desprolyl-enalaprilat co-folding with ACE1.

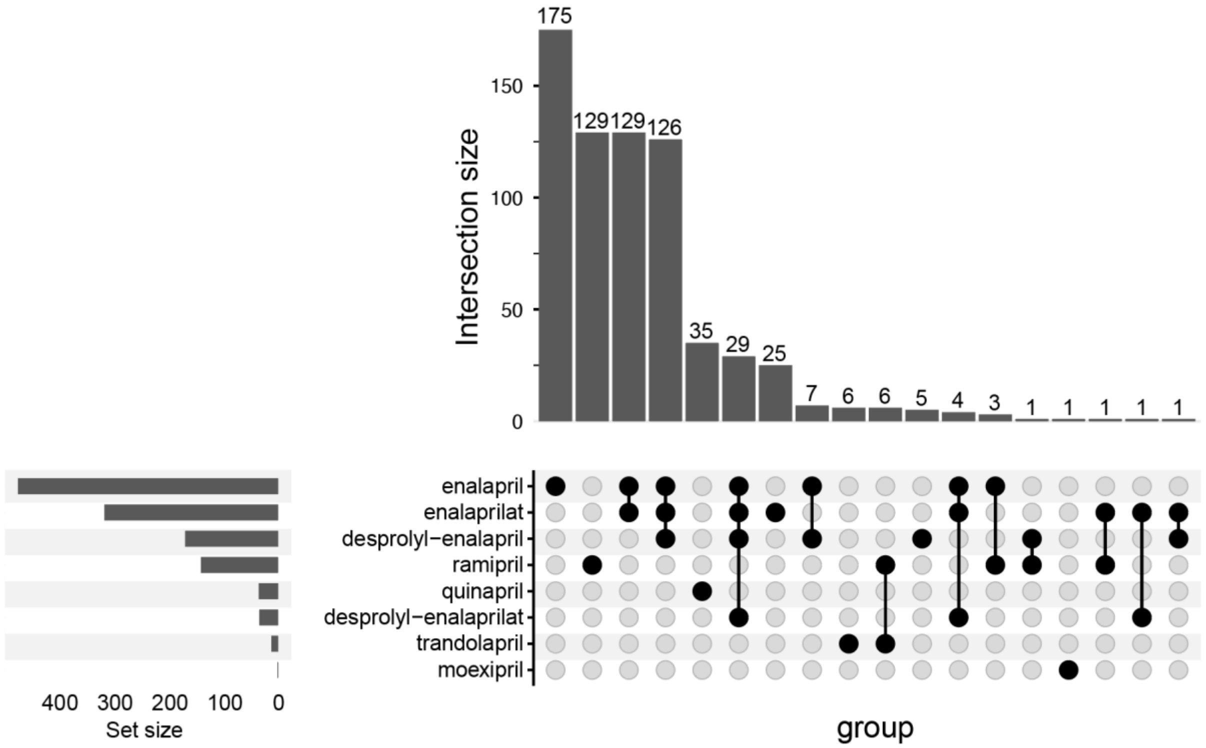


**Supplementary Figure 7 | Co-occurrence patterns of angiotensin-converting enzyme inhibitors and desprolyl metabolites in public datasets.** Upset plot showing co-occurrence frequencies for five angiotensin-converting enzyme inhibitors (enalapril, ramipril, quinapril, trandolapril, and moexipril) and the desprolyl metabolites (desprolyl-enalapril and desprolyl-enalaprilat) across public datasets of human, animal, and bacteria. Bars indicate the number of samples in which the compounds are detected, nodes denote the presence/absence for all intersections. Set size bars summarize the total of occurrence for each compound.
